# Sugar-rich larval diet promotes lower adult pathogen load and higher survival after infection in a polyphagous fly

**DOI:** 10.1101/2021.12.16.473033

**Authors:** Hue Dinh, Ida Lundbäck, Sheemal Kumar, Anh The Than, Juliano Morimoto, Fleur Ponton

## Abstract

Nutrition is a central factor influencing immunity and resistance to infection, but the extent to which nutrition during development affects adult responses to infections is poorly understood. Our study investigated how the nutritional composition of the larval diet affects the survival, pathogen load, and food intake of adult fruit flies, *Bactrocera tryoni*, after bacterial septic infection. We found a sex-specific effect of larval diet composition on survival post-infection: survival rate was higher and bacterial load was lower for infected females fed sugar-rich larval diet compared with females fed protein-rich larval diet, an effect that was absent in males. Both males and females were heavier when fed a balanced larval diet compared to protein- or sugar-rich diet, while body lipid reserves were higher in the sugar-rich larval diet compared with other diets. Body protein reserve was lower for sugar-rich larval diets compared to other diets in males, but not females. Both females and males shifted their nutrient intake to ingest a sugar-rich diet when infected compared with sham-infected flies without any effect of the larval diet, suggesting that sugar-rich diets can be beneficial to fight off bacterial infection. Overall, our findings show that nutrition during early life can shape individual fitness in adulthood.

**Summary statement:** Developmental conditions influence adulthood. Here we showed that when the larval diet is rich in sugar, resistance to infection is increased in females at adulthood, in a polyphagous fly.

## 1. Introduction

Environmental conditions during development can influence many aspects of adult phenotype and fitness. In humans, under- or over-nutrition at the foetal stage can increase predisposition to metabolic disease at the adult stage (see review in [1]). In birds, when conditions are unfavourable during early development, growth and adult immunity are affected ([2–6], but also see [7,8]). In holometabolous insects, the nutritional resources acquired at the larval stage are crucial for survival during metamorphosis [9,10]. When faced with food restrictions and unbalanced diets at the larval stage, insects show delayed development [11–16], lower adult body size [12,14,16–20], and decreased lifespan [21,22] (but see [11] where *Drosophila* adult life span is increased when fed a protein-restricted diet during larval stage). Adult reproductive performance is also affected by larval diet restriction with lower courtship level, lower number of mating, lower investment in reproductive organs, and a decrease in the total offspring production compared to individuals that were fed *ad libitum* at larval stage [14, 16, 17, 20, 23–26].

Nutrition during development affects not only life-history traits but also resistance to infection. In insects, food shortage at the larval stage strongly reduces immune activity, as observed in adult damselflies, *Lestes viridis* [27]. Larval food quality (i.e., yeast-to-sugar ratio) also influences the expression of antimicrobial peptide genes (Diptericin A and Metchnikowin) in adult *Drosophila*, with an increase in expression when the yeast-to-sugar ratio in the larval diet was increased [28]. In the cotton leafworm, *Spodoptera littoralis*, a change in the larval diet affects lytic and phenoloxidase activity without any evidence of change in immune gene expression [29]. The effects of larval diet on immune-challenged adults have recently been described in female *Anopheles coluzzii* with an effect on the prevalence and intensity of *Plasmodium berghei* infection [30]. However, it remains unclear which aspect of the larval diet (quality or quantity) induces differences in pathogen load and survival rate of the infected individuals. Interestingly, in hemimetabolous insects, Kelly et al. (2013) found a sex-specific effect of juvenile diet on the survival of adult crickets, *Gryllus texensis*, when infected with the pathogenic bacterium, *Serratia marcescens*, using defined diets (i.e., low- or high-protein diets) [31]. However, crickets were fed the same experimental diet at both juvenile and adult stages, and, therefore, it is difficult to decipher between the effects of juvenile and adult diet on adult resistance in this system.

Studies on the effects of developmental diet on adult immunity and resistance to infection, especially when adults are immune challenged, are still scarce or have been only partial for three main reasons. First, diet manipulations during development have focused on the quantity of food available, and there have been very few studies investigating how diet composition might affect adult resistance. The nutritional environment is likely to vary not only on the quantity of food available but also in the quality of food sources with nutritional imbalances. Second, the nutritional requirements of immune responses can be different between sexes [32]. For instance, encapsulation ability increases with the intake of both protein and sugar in females of the decorated crickets, whereas male encapsulation ability only increases with protein intake [33]. Hence, it remains to be tested if pathogen resistance in both adult males and females is affected similarly by the juvenile diet. Lastly, individuals might compensate for unfavourable juvenile nutritional conditions by modifying their diet at the adult stage. Exploring the effects of variations in the quality of juvenile diet on adult nutritional responses and body energetic reserves would give us insights into the extent to which nutritional conditions early in life modulate adult physiology and fitness.

Here, we manipulated the dietary macronutrient ratio (yeast-to-sugar ratio, YS) during the larval stage of the holometabolous fruit fly *Bactrocera tryoni*. We investigated the effects of larval diet manipulation on (i) developmental traits (i.e., percentage of egg hatching, percentage of pupation, percentage of emergence and developmental time); (ii) adult physiological traits (i.e., total body weight, body lipid, and protein); and (iii) adult response to a septic infection with the pathogenic bacterium *S. marcescens* (i.e., bacterial load, survival, and food intake). Our findings provide new insights into how environmental experience during the larval stage influences a broad range of life-history traits as well as the outcome of septic infection in adulthood.

## 2. Materials and Methods

### Fly stock

Fly stock was maintained on a gel-based diet at the larval stage [34], and a 1:3 ratio of hydrolysed yeast (MP Biomedicals Cat. no 02103304) to sugar (CSR® White Sugar) (YS) was provided separately at the adult stage. Flies were reared in a controlled environment room under the conditions of 25^0^C and 65% humidity, with a 12-hour light/dark cycle at Macquarie University ARC Centre for Fruit Fly Biosecurity Innovation (North Ryde, NSW, Australia). Eggs were collected from the fly stock colony for 2 h using an ovipositional device that consisted of a plastic bottle with numerous puncture holes and filled with 30 mL of water to maintain humidity. The collected eggs were used to assess the effects of developmental diet on development and adult traits.

### Diet preparation

Three larval diets varying in the yeast-to-sugar ratio (YS) were prepared (listed in Table S1). The standard diet is considered optimized for larval development, and it has been used routinely to rear *B. tryoni* (YS 1.67:1) [34]. We manipulated the relative amount of yeast and sugar [34] to generate unbalanced diets, including a “protein-rich diet” (YS 5:1) and a “sugar-rich diet” (YS 1:3.4). These diets have been found to modulate the development and adult life-history traits of *B. tryoni* flies [35]. The yielding percentages of protein (w/w (Y + S)) in the three substrates were 70% (YS 5:1), 43% (YS 1.67:1), and 12% (YS 1:3.4). All ingredients were mixed into warm water and the final volume (250 mL) was achieved by adding distilled water. Citric acid was added to adjust the pH of the diet solution to 3.5 at room temperature. To assess developmental traits, diet plates were prepared by pouring 25 mL of larval gel diet into 100 mm. When we needed to rear a large number of larvae, 150ml of diet was poured into plastic trays (17.5 cm long, 12 cm wide, 4 cm deep).

### Development traits

Groups of 100 eggs were transferred to a black filter paper previously soaked in distilled water and placed onto the diet plates. The plates were then covered with their lids and kept under controlled laboratory conditions during larval development. Nine replicates per larval diet treatment were performed simultaneously (i.e., 9 plates). The number of unhatched eggs were counted 4 days post-seeding under a stereomicroscope, and the black filter paper and unhatched eggs removed from the diet plates. Lids of the diet plates were opened seven days post-seeding; plates were then placed on 50 mL of autoclaved fine vermiculite to allow larvae to jump outside the plates and pupate. The total number of pupae was then recorded for each plate, and pupae was placed into partially netted 12.5 litre plastic cages for emergence (9 replicates). The number of pupae that did not emerge was recorded over four days.

### Adult traits

Forty ml of eggs (∼600 eggs) was seeded into 150 ml diet to achieve the same density as we had in the developmental experiment (100 eggs per 25ml diet). Eggs were allowed to develop until the adult stage. One-day-old adults (i.e., collected one day after eclosion) were used for the different measurements.

#### Adult dry body weight

Flies were collected and stored at -20^0^C. Carcasses were dried at 55^0^C for 48 h (Binder drying oven). Dry weight was measured using a microbalance (Sartorius, accuracy ±0.001mg) for 30 individual flies of each sex per diet treatment.

#### Adult body reserves

Body lipid reserves were extracted in three, 24-h changes of chloroform as previously described [36]. At the end of the third chloroform wash, lipid-free bodies were re-dried and re-weighed to calculate lipid content. We performed 15 replicates (i.e., 15 individual flies of each sex) per diet treatment.

#### Adult body protein reserves

After lipid extraction, fly bodies were crushed in 300 µl 0.1M NaOH and centrifuged at 8000 rpm for 30 sec. 100 µl of supernatant was collected in new Eppendorf tubes and diluted 1:10 time. Five µl of the diluted solutions were transferred to 96-well plates and allowed to react with 200 µl of Bradford reagent (Sigma-Aldrich). Plates were incubated for 5 min at room temperature, and absorbance was measured at 595 nm using a spectrometer (Eppendorf). We ran 15 biological replicates (i.e., 15 individual flies of each sex) per diet treatment. Each sample was run in 3 technical replicates. The Bradford assay was calibrated using a standard curve generated from 6 different concentrations of IgG protein (Sigma-Aldrich) (0.2, 0.15, 0.1, 0.05, 0.025, and 0 µg/µl).

### Bacterial infection

*Serratia marcescens* (ATCC 13880, Thermo Scientific) was inoculated into 5 mL of sterile Nutrient Broth (Oxoid, CM0001) and incubated overnight (approximately 16 hrs) at 26^0^C with shaking at 200 rpm. The bacterial culture was centrifuged at 10,000g at 4^0^C for 2 min. The supernatant was discarded, and the bacterial pellet washed twice using 1X Phosphate Buffered Saline (PBS) (Sigma-Aldrich, Cat. No P4417) to remove any trace of the medium. The bacterial pellet was resuspended to a target concentration of OD_600_ = 0.025 in sterile PBS.

One day after adult eclosion, flies were cold anesthetized at -20^0^C for 2 minutes and placed on a Petri dish on a dry bath (Product code: MK20) at -10°C. Injections were performed using a 10µL syringe (NanoFil) connected to a microinjector (World Precise Instrument) with a delivery speed of 50 nL/sec. A volume of 0.2 µL of the bacterial solution, yielding a dose of approximately 1680 cells, was injected into the fly’s coxa of the third right leg. PBS-injected (i.e., sham-injured) flies were used as controls.

### Bacterial load

Bacterial load was measured in infected flies with females and males being individually crushed in 100 µL of PBS and serially diluted to 1:10 and 1:100. A volume of 10 µL from each dilution was plated onto Nutrient Agar supplemented with 30 µg/mL Tetracycline (Sigma) [36] and incubated at 26^0^C for 48 h. The bacterial load was measured 6 h, and 1, 2, and 4 days post-infection (PI) (10 cages per diet). We sampled 1 fly per replicate cage (i.e., 10 individual flies of each sex) for each diet treatment at each time point. *Serratia marcescens* was not present in our fly stock. This was checked by crushing individual flies (12 individual flies of each sex) in 100 μl PBS, and 25 μl of the solution was plated onto Nutrient Agar supplemented with 30 μg/mL Tetracycline, and incubated at 26 °C for 48 h. We did not observe any *S. marcescens* colonies.

### Survival after infection

One day after adult eclosion, adult flies (males and females) were injected with either PBS or live bacteria. Injected flies were then maintained in groups of 25 in 1.25-liter cages (10 cm × 10 cm × 12.5 cm) and provided with food and water *ad libitum*. Dead flies were counted and removed daily from the cages. We initially limited the experimental time to 4 days post-infection (PI), in which we measured bacterial load. The low mortality rate after 4 days PI led us to extend the timeframe of the survival experiment until 15 days PI. We ran 3 replicates (3 cages) per diet.

### Food intake

The method to measure and calculate food intake was previously described in [36]. Briefly, flies were housed individually and allowed to self-select between a sugar (CSR® White Sugar) and a yeast (MP Biomedicals Cat. No. 02103304) solution. Sugar and yeast were provided separately in two 30 µL capillaries at a final concentration of 160 g/L. The hydrolysed yeast used in this study was the only source of protein available to the flies, containing approximately 62.1% protein and 1% sugar. Final macronutrient (i.e., protein and sugar) intakes (µg) were calculated based on these values.

### Statistical analyses

Statistical analyses were performed using R [37], and graphs were done using BM SPSS Statistics 25.0. We fitted Generalized Linear Models (GLM) with quasibinomial distribution to analyse the proportion of egg hatching, pupation, emergence, and body reserves. We fitted Generalized Linear Models (GLM) with Gaussian distribution to analyse the dry body weight and bacterial load (log-transformed) of infected flies. To analyse the survival data, we could not use a Cox regression because the proportional-hazards assumption was not respected (p global=0.017). We, therefore, analysed the percentage of flies that died 4 and 15 days PI using GLM with a quasibinomial distribution. Because only a very small number of PBS-injected flies died during the course of the experiment (15 days), the analysis was performed only for infected flies across diets.

## 3. Results

### Effects of larval diet on developmental traits

Larval diet did not influence the percentage of egg hatching (GLM, F_2,24_ = 0.01, P = 0.905), percentage of pupation (GLM, F_2,23_ = 1.940, P = 0.166) and percentage of emergence (GLM, F_2,23_ = 1.111, P = 0.346). The percentage of egg hatching and percentage of pupation were around 90% in all larval diet treatments. The percentage of emergence observed across larval diets was around 98%. Larval diet had however a significant effect on the egg-to-adult developmental time (GLM, F_2,23_ = 33.896, P < 0.001). As expected, developmental time was longer for the larvae fed the sugar-rich larval diet (20.13 ± 0.641 days) compared to those fed the balanced (18.33 ± 0.500 days) and protein-rich larval diets (18.22 ± 0.441 days).

### Effects of larval diet on adult body weight, total body lipid, and body protein

The dry body weight was significantly influenced by larval diet and sex (GLM; Sex: F_(1,81)_ = 32.59, P<0.001; Larval diet: F_2,82_ = 4.64, P = 0.012; Larval diet x Sex: F_2,79_ = 3.73, P = 0.078). Adults flies reared in the standard diet at the larval stage had a higher body weight relative to those reared in either the protein-rich or sugar-rich larval diet (Fig. 1a). Additionally, adult body weight was higher in females compared to males (Fig. 1A). The percentage of body lipid reserves was significantly influenced by larval diet for both females and males (GLM; Sex: F_1,81_ = 0.111, P = 0.739; Larval diet: F_2,82_ = 26.1868, P < 0.001; Larval diet x Sex: F_2,79_ = 1.165, P = 0.312). Body lipid reserves were greater in adult flies from the sugar-rich larval diet compared to flies from the balanced and protein-rich diets (Fig. 1b). The interaction between sex and larval diet composition significantly influenced the percentage of body protein reserves (GLM; Sex: F_1,81_ = 25.522, P < 0.001; Larval diet: F_2,82_ = 182.757, P < 0.001; Larval diet x Sex: F_2,79_ = 14.928, P < 0.001). The body protein reserves of males from the sugar-rich larval diet were lower compared to those of males from the protein-rich and balanced larval diets (Fig. 1c). We did not detect any effect of the larval diet on females’ body protein reserves (Fig. 1c).

**Figure 1.**
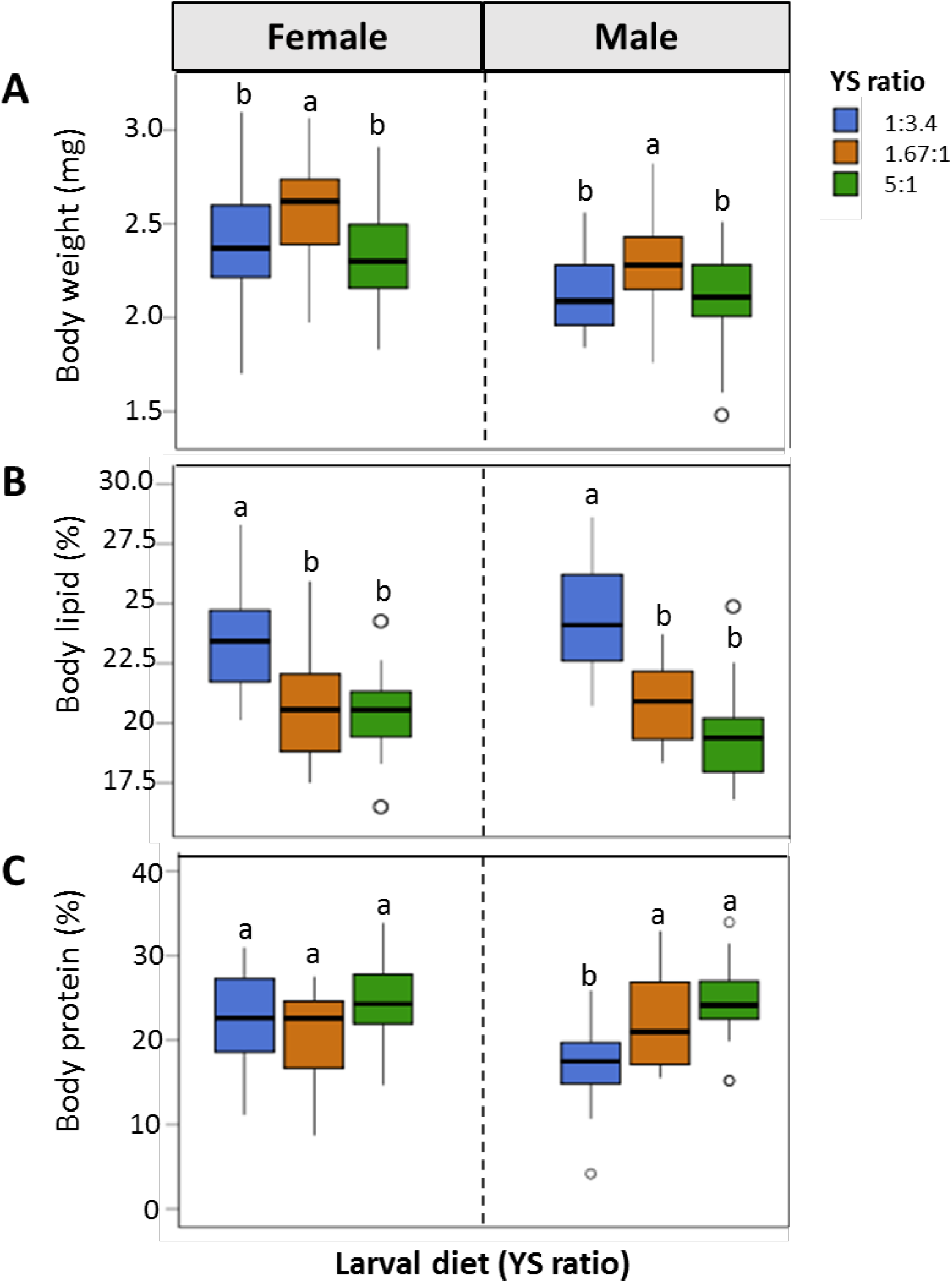
Effect of larval diet on adult body weight, lipid and protein body reserves. Body weight (A), lipid body reserves (B) and protein body reserves (C) were measured in male and female flies fed three larval diets varying in the yeast-to-sugar ratio (YS ratio). Different letters indicate significant differences between larval diet treatments, assessed by SNK test at P<0.05.

### Effects of larval diet on bacterial loads of adult flies

The two-way interaction between larval diet and time influenced the bacterial load of infected flies (P < 0.001; Table 1). Bacterial loads were comparable between larval diets at 6 and 48 h PI (Fig. 2A). At 24 h PI, bacterial load tended to be higher in flies fed the protein-, and sugar-rich larval diets relative to flies fed the balanced diet (Fig. 2a). At 96 h PI, the bacterial load of flies fed the yeast-rich diet was greater compared to that of flies fed either the balanced or sugar-rich larval diet (Fig. 2A).

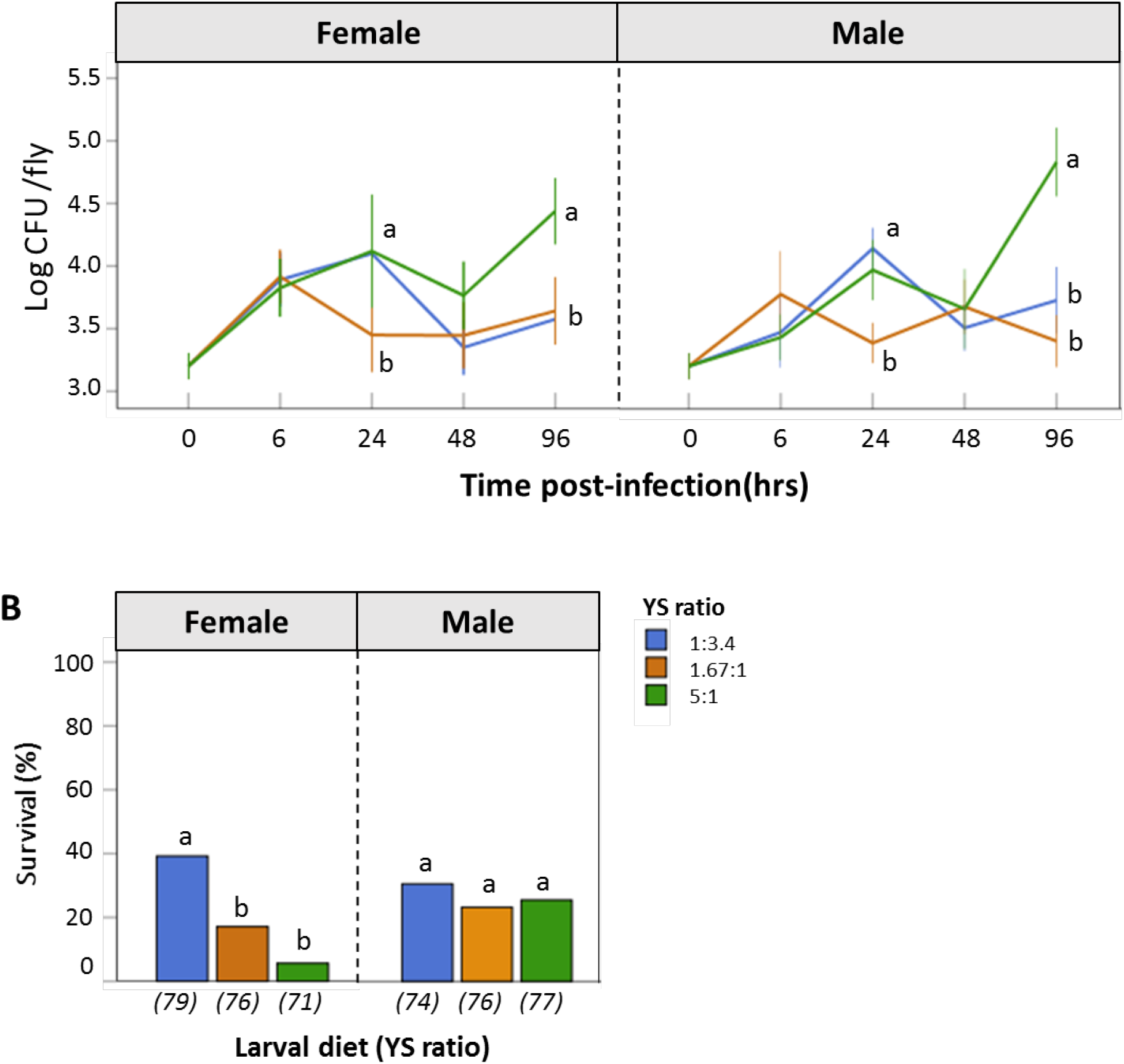
(A) Effect of larval diets on bacterial loads measured at 0, 6, 24, 48 and 96 h post-infection when fed three larval diets varying in the yeast-to-sugar ratio (YS ratio). Blue line - YS 1:3.4 (sugar-rich larval diet); Orange line - YS 1.67:1 (balanced larval diet); Green line - YS 5:1 (protein-rich larval diet). (B) Effects of larval diets on the survival rate of infected females and males at 15 days post-infection. Larval diet varied in the yeast-to-sugar ratio (YS ratio). Numbers in parentheses below the bars indicate number of flies in each treatment. In both (A) and (B), different letters indicate significant differences between larval diets, assessed by SNK test at P<0.05.

**Table 1.** GLM outcome for the effects of larval diet, sex and time on bacterial load (log-transformed).

| Response variable | factor | df | Resid. df | F-value | P-value |
| --- | --- | --- | --- | --- | --- |
| Bacterial load | Larval diet | 2 | 295 | 14.145 | **<0.001** |
|  | Sex | 1 | 297 | 0.506 | 0.477 |
|  | Time | 1 | 298 | 36.757 | **<0.001** |
|  | Larval diet:Sex | 2 | 290 | 0.049 | 0.952 |
|  | Larval diet:Time | 2 | 292 | 24.535 | **<0.001** |
|  | Sex:Time | 1 | 294 | 3.505 | 0.062 |
|  | Diet:Sex:Time | 2 | 288 | 1.877 | 0.155 |

### Effects of larval diet on survival of infected flies

At 4 days PI, the survival of infected flies was not affected by larval diet or sex (Larval diet: F_2, 450_ = 1.421, P = 0.241; Sex: F_1, 449_ = 0.291, P = 0.589; Larval diet x Sex: F_2, 447_ = 2.500, P = 0.083). At 15 days PI, however, we observed a significant effect of the interaction between larval diet and sex (Larval diet: F_2, 450_ = 2.949, P = 0.053; Sex: F_1, 449_=0.098, P=0.754; Larval diet x Sex: F_2, 447_ = 4.600, P = 0.010). Survival of infected females from the sugar-rich larval diet was significantly higher compared to those from the balanced and protein-rich larval diets (Fig. 2B); however, we did not detect any effects of larval diet on the survival of infected males (Fig. 2B).

### Effects of larval diet on the nutritional choice of immune-challenged adult male and female flies

The ingested macronutrient ratio (protein-to-sugar ratio, PS ratio) was influenced by treatment and sex (P < 0.001; Table 2). Infected flies ingested a lower PS ratio [i.e., diet richer in sugar, PS ∼0.302 (1:3.3)] compared to PBS-injected flies [i.e., PS ∼0.537 (1:1.8)] (Fig. 3). Females ingested a diet that was slightly richer in protein than males [PS females∼0.430 ± 0.203 (1:2.3); PS males∼0.373 ± 0.180 (1:2.7)]. The amount of protein ingested after 4 days was influenced by the interaction between larval diet and injection treatment (P = 0.021, Table 2). There was a trend for flies from the sugar-rich larval diet to ingest less protein than the individuals from the 2 other larval diets (Fig. 3). This trend was more marked in PBS-injected individuals than in bacteria-injected ones, certainly, because infected ones ingested less food (Fig. 3). The total quantity of sugar ingested after 4 days was slightly influenced by the interaction between larval diet and sex (P = 0.050, Table 2). As observed for protein intake, individuals from the sugar-rich larval diet tended to ingest less sugar (Fig. 3). In females, sugar intake tended to increase with the protein content of the larval diet (Fig. 3); particularly, infected females reared on the protein-rich diet ingested the highest quantity of sugar (Fig. 3). This trend was also observed in males but was less clear (Fig. 3). Males tended to ingest less food than females (Fig. 3).

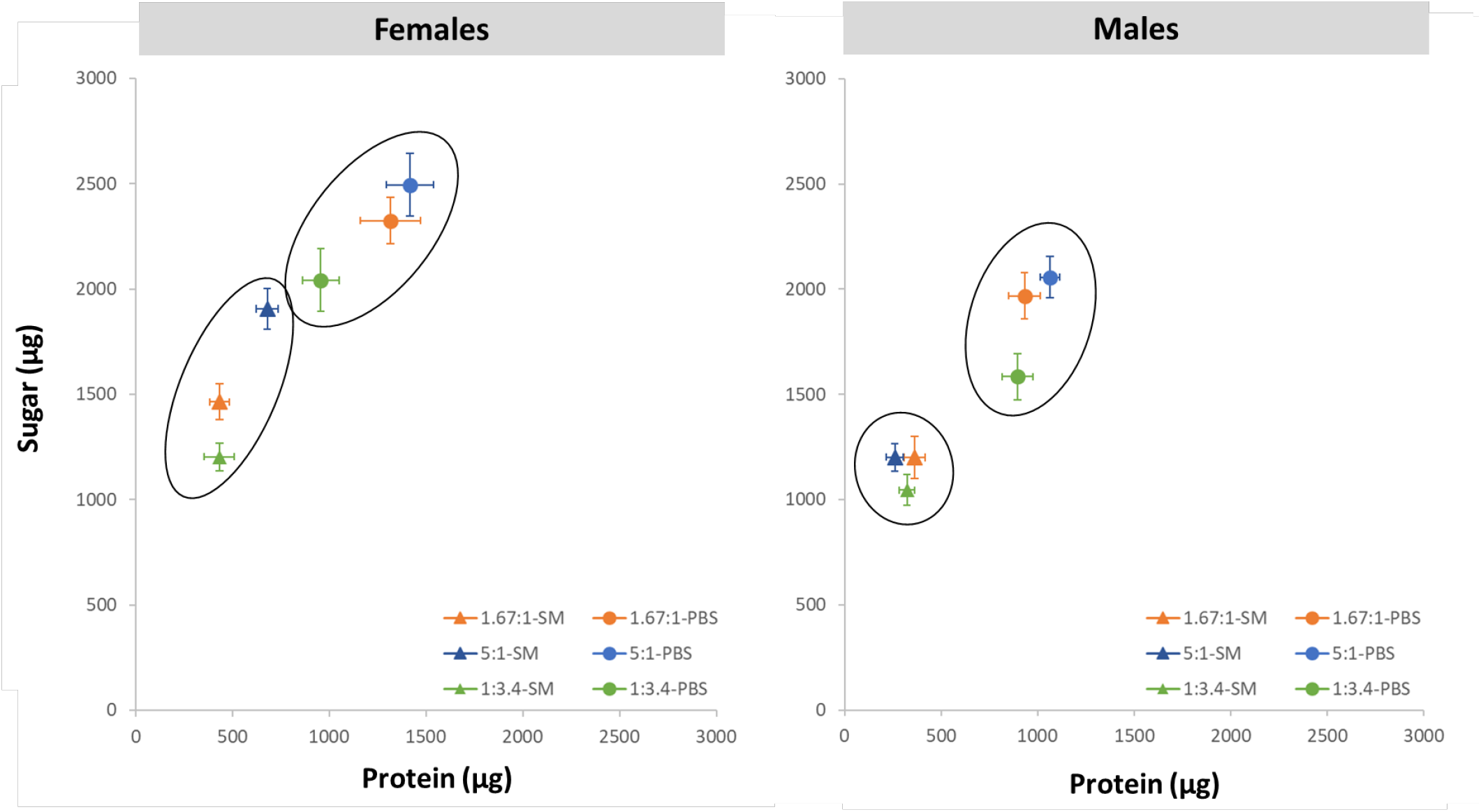

**Table 2.** GLM outcome for the effects of larval diet, sex and injection treatment on feeding behaviour at adult stage.

| Response variable | factor | df | Resid. df | F-value | P-value |
| --- | --- | --- | --- | --- | --- |
| Protein-to-sugar ratio | Larval diet | 2 | 160 | 0.078 | 0.924 |
|  | Sex | 1 | 159 | 5.765 | **0.017** |
|  | Injection | 1 | 158 | 92.252 | **<0.001** |
|  | Larval diet:Sex | 2 | 156 | 2.121 | 0.123 |
|  | Larval diet:Injection | 2 | 154 | 0.637 | 0.530 |
|  | Sex:Injection | 1 | 153 | 0.709 | 0.401 |
|  | Larval diet:Sex:Injection | 2 | 151 | 1.463 | 0.235 |
| Total protein intake | Larval diet | 2 | 160 | 7.204 | **0.001** |
|  | Sex | 1 | 159 | 234.814 | **<0.001** |
|  | Injection | 1 | 158 | 26.814 | **<0.001** |
|  | Larval diet:Sex | 2 | 156 | 2.628 | 0.075 |
|  | Larval diet:Injection | 2 | 154 | 3.969 | **0.021** |
|  | Sex:Injection | 1 | 153 | 0.556 | 0.457 |
|  | Larval diet:Sex:Injection | 2 | 151 | 1.928 | 0.149 |
| Total sugar intake | Larval diet | 2 | 160 | 21.017 | **<0.001** |
|  | Sex | 1 | 159 | 53.670 | **<0.001** |
|  | Injection | 1 | 158 | 149.709 | **<0.001** |
|  | Larval diet:Sex | 2 | 156 | 3.054 | **0.050** |
|  | Larval diet:Injection | 2 | 154 | 0.393 | 0.676 |
|  | Sex:Injection | 1 | 153 | 0.055 | 0.815 |
|  | Larval diet:Sex:Injection | 2 | 151 | 1.911 | 0.151 |

## 4. Discussion

We examined the effects of the macronutrient composition of larval diet on adult resistance to infection as well as on some developmental and physiological traits. When adult flies were challenged with *S. marcescens*, we observed a higher bacterial load in both males and females fed a protein-rich larval diet; however, only females showed a lower survival rate after infection. This might be partly explained by the result that body lipid reserves were greater in adult flies from the sugar-rich larval diet compared to flies from the balanced and protein-rich diets, the body protein reserves of males from the sugar-rich larval diet were also lower compared to those of males from the protein-rich and balanced larval diets. However, there was no effect of the larval diet on females’ body protein reserves. The larval diet also influenced adult feeding choice, with flies from the sugar-rich larval diet ingesting slightly less protein than the individuals from the 2 other larval diets.

### Larval diet influences adult pathogen resistance and macronutrient intake following infection with the bacterium S. marcescens

The bacterial load of infected flies was influenced by the nutritional conditions experienced at the larval stage. This might be due to two reasons. First, pathogens require energy for growing, and thus, the allocation of within-host energy reserves is essential to this process [38–42]. Here, we found that the total body reserves of protein and/or lipid were modulated by larval diets. Second, early-life nutrition affects host immune responses at later developmental stages, which might modulate the number of pathogenic cells in the host. In mosquitoes and *Drosophila*, poor larval conditions (i.e., starvation or protein-restriction) have been shown to alter the expression of adult immune-related genes [28, 43].

Despite differences in the bacterial load between larval diet treatments, the survival of infected flies was similar during the first 4 days post-infection. Infected flies might only start dying when the number of pathogenic bacteria reaches a certain level defined as “bacterial load upon death” [44], and this level may not have been reached only a few days after the infection. At 15 days PI, however, infected females that were reared on the sugar-rich larval diet survived at a greater rate compared to those kept on the protein-rich larval diet. Several hypotheses can explain these results. First, a lower bacterial load at the early stage of infection might have slowed down the time required to reach the “bacterial load upon death” in infected females fed the sugar-rich larval diet. Second, it has been previously shown that infected flies reduce total food intake and shift diet choice towards a sugar-rich diet which promote their survival after infection [36]. We also found that infected adult flies ingested a lower PS ratio compared to PBS-injected flies (similar result as in [45]). Further, flies from the sugar-rich larval diet tended to ingest less protein than the individuals from the 2 other larval diets, which might provide them better resistance to infection for the positive effects of anorexia on host defence. Third, female flies fed the sugar-rich larval diet might have invested more in immunity at the expense of other life-history traits. This is supported by studies in birds and insects showing the negative correlation between immune function and developmental time [46–47]. For instance, female moths reared on a sugar-rich larval diet allocate a lower proportion of their mass to the development of their ovaries (i.e., invest less on reproduction) compared to those on protein-rich diets [14]. Hence, female flies fed the sugar-rich larval diet potentially prioritize their immunity over developmental time (see also [48] for a similar discussion). Also, fat content serves as a crude estimate of the size of the fat body, the major immune responsive tissue in insects [49]. Because body lipid reserves were higher in flies fed the sugar-rich larval diet, it is possible that their immune system was more efficient at fighting the infection. Further explorations of the immune status, reproductive output, and longevity of females kept on the different larval diets would provide insights into the effects of the juvenile nutritional environment on resource allocation and potential trade-offs between immune traits and other life-history traits.

Early-life environment, including nutrition, can be used to predict future adult environment, and individuals can develop proper behaviours to respond to environmental challenges in later developmental stages [50]. The higher survival rate of infected females on a sugar-rich larval diet might suggest that unbalanced diets at the larval stage can act as a cue for higher disease risk in adulthood, and flies on this diet might invest more in defence (see also [30, 31, 51–53]). This does not, however, explain the sex-dependent effects in our results.

Unlike what we observed in female flies, the survival rates of infected adult males were comparable between the larval diet treatments despite a difference in bacterial load. While it is difficult at this stage to explain why the effect of larval diet on survival rate is sex-specific, previous studies have shown that the diet composition can influence immunity differently in males and females. For instance, in fruit flies and crickets, while both protein and sugar intakes affect phenoloxidase (PO) activity and encapsulation ability in females, only protein intake influences these immune traits in males [32,33]. Also, the magnitude of the effects of larval diet composition on PO activity and nitric oxide production in adults can be different between male and female mosquitoes, *Aedes aegypti*, with a stronger effect in females [54]. In parallel, a larval diet can influence differently other adult traits in both sexes. In the butterfly *Melitaea cinxia*, larval food stress negatively affects the reproductive output of females-but not males [55]. Also, developing on a high yeast diet only benefits life span of female *Drosophila* [56]. While there are evidences of sex-specific effects of diet on adult traits, the physiological mechanisms at the basis of these differences remain to be investigated.

### Effect of larval diet on development traits

The percentage of pupation and percentage of emergence were similar between flies from the different larval diet treatments, suggesting that the unbalanced larval diets chosen here did not affect larval ability to survive metamorphosis. However, we found a significant effect of the larval diet on egg-to-adult development time, and interestingly, this was only observed in larvae fed the sugar-rich diet. This result is in accordance with previous observations in the forest tent caterpillar, *Malacosoma disstria* [14,57]. The negative effect of the sugar-rich larval diet on development time is likely caused by the low protein level. Insect growth and metamorphosis are controlled by the insulin/target of rapamycin (TOR) signalling pathways [58–60], which are triggered by high levels of amino acids [61,62]. Indeed, inhibition of the amino acid transporter gene has been shown to result in a lengthened development time [63]. Measuring the insulin/TOR activity in larvae fed the experimental diets would give insights into their metabolic state and deepen our understanding of the links between low-protein feeding and delayed developmental time.

## 5. Conclusion

The present study highlights the sex-specific effect of the larval diet composition on the survival of adult fruit flies after infection. Protein-rich larval diet promoted higher bacterial load and lower survival in female flies. The profound effects of larval diet on the developmental and physiological traits of adults were also demonstrated. Better understanding the carry-over effects of environmental conditions experienced in early life on individuals life-history traits and population dynamic is a central question in ecology [64]. Answers to this question can further assist the protection of endangered species, especially in the context of dramatic environmental changes that potentially lead to decreases in food availability and changes in food composition as well as introductions of infectious disease to wildlife populations [65,66].

## Acknowledgments

We thank Prof. Phillip Taylor for laboratory support and fruitful discussions regarding the data analyses.

## Author contributions

H.D. designed and carried out the experiments and performed the statistical analyses. I.L. carried out the experiments, collected and analysed the data. S.K. carried out the experiments and analysed the data. A.T.T. helped in setting up the experiments. J.M. performed the statistical analyses. F.P. conceived and designed the experiments and performed the statistical analyses. All authors contributed to the writing of the manuscript and gave final approval for publication.

## Data availability

Data will be deposited in Dryad after acceptance for publication of the paper.

## References

[1] Gluckman PD, Hanson MA, Cooper C, Thornburg KL. Effect of in utero and early-life conditions on adult health and disease. N Engl J Med 2008;359:61. doi:10.1056/NEJMra0708473.

[2] Butler MW, McGraw KJ. Past or present? Relative contributions of developmental and adult conditions to adult immune function and coloration in mallard ducks (Anas platyrhynchos). J Comp Physiol B Biochem Syst Environ Physiol 2011;181:551–63. doi:10.1007/s00360-010-0529.

[3] Naguib M, Nemitz A, Gil D. Maternal developmental stress reduces reproductive success of female offspring in zebra finches. Proc R Soc B Biol Sci 2006;273:1901–5. doi:10.1098/rspb.2006.3526.

[4] Naguib M, Riebel K, Marzal A, Gil D. Nestling immunocompetence and testosterone covary with brood size in a songbird. Proc R Soc B Biol Sci 2004;271:833–8. doi:10.1098/rspb.2003.2673.

[5] Tella JL, Forero MG, Bertellotti M, Donázar JA, Blanco G, Ceballos O. Offspring body condition and immunocompetence are negatively affected by high breeding densities in a colonial seabird: A multiscale approach. Proc R Soc B Biol Sci 2001;268:1455–61. doi:10.1098/rspb.2001.1688.

[6] Krause ET, Honarmand M, Wetzel J, Naguib M. Early fasting is long-lasting: Differences in early nutritional conditions reappear under stressful conditions in adult female zebra finches. PLoS One 2009;4:e5015. doi:10.1371/journal.pone.0005015.

[7] Råberg L, Stjernman M. Natural selection on immune responsiveness in blue tits Parus caeruleus. Evolution (N Y) 2003;57:1670. doi:10.1554/02-417.

[8] Tschirren B, Rutstein AN, Postma E, Mariette M, Griffith SC. Short- and long-term consequences of early developmental conditions: A case study on wild and domesticated zebra finches. J Evol Biol 2009;22:387–95. doi:10.1111/j.1420-9101.2008.01656.x.

[9] Arrese EL, Soulages JL. Insect Fat Body: Energy, metabolism, and regulation. Annu Rev Entomol 2010;55:207–25. doi:10.1146/annurev-ento-112408-085356.

[10] Mirth CK, Riddiford LM. Size assessment and growth control: How adult size is determined in insects. BioEssays 2007;29:344–55. doi:10.1002/bies.20552.

[11] May CM, Doroszuk A, Zwaan BJ. The effect of developmental nutrition on life span and fecundity depends on the adult reproductive environment in Drosophila melanogaster. Ecol Evol 2015;5:1156–68. doi:10.1002/ece3.1389.

[12] Zwaan BJ, Bijlsma R, Hoekstra RF. On the developmental theory of ageing. I. starvation resistance and longevity in Drosophila melanogaster in relation to pre-adult breeding conditions. Heredity (Edinb) 1991;66:29–39. doi:10.1038/hdy.1991.4.

[13] Telang A, Frame L, Brown MR. Larval feeding duration affects ecdysteroid levels and nutritional reserves regulating pupal commitment in the yellow fever mosquito Aedes aegypti (Diptera: Culicidae). J Exp Biol 2007;210:854–64. doi:10.1242/jeb.02715.

[14] Colasurdo N, Gelinas Y, Despland E. Larval nutrition affects life history traits in a capital breeding moth. J Exp Biol 2009;212:1794–800. doi:10.1242/jeb.027417.

[15] Roeder KA, Behmer ST. Lifetime consequences of food protein-carbohydrate content for an insect herbivore. Funct Ecol 2014;28:1135–43. doi:10.1111/1365-2435.12262.

[16] Xie J, De Clercq P, Pan C, Li H, Zhang Y, Pang H. Larval nutrition-induced plasticity affects reproduction and gene expression of the ladybeetle, Cryptolaemus montrouzieri. BMC Evol Biol 2015;15:276. doi:10.1186/s12862-015-0549-0.

[17] Morimoto J, Ponton F, Tychsen I, Cassar J, Wigby S. Interactions between the developmental and adult social environments mediate group dynamics and offspring traits in Drosophila melanogaster. Sci Rep 2017;7:3574. doi:10.1038/s41598-017-03505-2.

[18] Morimoto J, Nguyen B, Dinh H, Than AT, Taylor PW, Ponton F. Crowded developmental environment promotes adult sex-specific nutrient consumption in a polyphagous fly. Front Zool 2019;16:4. doi:10.1186/s12983-019-0302-4.

[19] Boggs CL, Freeman KD. Larval food limitation in butterflies: effects on adult resource allocation and fitness. Oecologia 2005;144:353–61. doi:10.1007/s00442-005-0076-6.

[20] Wang Y, Kaftanoglu O, Brent CS, Page RE, Amdam G V. Starvation stress during larval development facilitates an adaptive response in adult worker honey bees (Apis mellifera L.). J Exp Biol 2016;219:949–59. doi:10.1242/jeb.130435.

[21] Bauerfeind SS, Fischer K. Effects of larval starvation and adult diet-derived amino acids on reproduction in a fruit-feeding butterfly. Entomol Exp Appl 2009;130:229–37. doi:10.1111/j.1570-7458.2008.00814.x.

[22] Runagall-Mcnaull A, Bonduriansky R, Crean AJ. Dietary protein and lifespan across the metamorphic boundary: Protein-restricted larvae develop into short-lived adults. Sci Rep 2015;5:11783. doi:10.1038/srep11783.

[23] Morimoto J, Pizzari T, Wigby S. Developmental environment effects on sexual selection in male and female Drosophila melanogaster. PLoS One 2016;11:e0154468. doi:10.1371/journal.pone.0154468.

[24] Kaspi R, Mossinson S, Drezner T, Kamensky B, Yuval B. Effects of larval diet on development rates and reproductive maturation of male and female Mediterranean fruit flies. Physiol Entomol 2002;27:29–38. doi:10.1046/j.1365-3032.2001.00264.x.

[25] Dmitriew C, Rowe L. The effects of larval nutrition on reproductive performance in a food-limited adult environment. PLoS One 2011;6:e17399. doi:10.1371/journal.pone.0017399.

[26] Yan J., Kibech R, Stone C.M. Differential effects of larval and adult nutrition on female survival, fecundity, and size of the yellow fever mosquito, Aedes aegypti. Frontiers in Zoology 2021 18(1):10. doi.org/10.1186/s12983-021-00395-z

[27] Rolff, Van de Meutter, Stoks. Time constraints decouple age and size at maturity and physiological traits. Am Nat 2017;164:559. doi:10.2307/3473403.

[28] Fellous S, Lazzaro BP. Larval food quality affects adult (but not larval) immune gene expression independent of effects on general condition. Mol Ecol 2010;19:1462–8. doi:10.1111/j.1365-294X.2010.04567.x.

[29] Cotter SC, Reavey CE, Tummala Y, Randall JL, Holdbrook R, Ponton F, Simpson SJ, Smith JA, Wilson F. Diet modulates the relationship between immune gene expression and functional immune responses. Ins Biochem Mol Biol 2019;109:128–141.

[30] Linenberg I, Christophides GK, Gendrin M. Larval diet affects mosquito development and permissiveness to Plasmodium infection. Sci Rep 2016;6:38230. doi:10.1038/srep38230.

[31] Kelly CD, Tawes BR. Sex-specific effect of juvenile diet on adult disease resistance in a field cricket. PLoS One 2013;8:e61301. doi:10.1371/journal.pone.0061301.

[32] Fanson BG, Fanson K V., Taylor PW. Sex differences in insect immune function: A consequence of diet choice? Evol Ecol 2013;27:937–47. doi:10.1007/s10682-013-9638-y.

[33] Rapkin J, Jensen K, Archer CR, House CM, Sakaluk SK, Castillo E Del, et al. The Geometry of nutrient space–based life-history trade-offs: sex-specific effects of macronutrient intake on the trade-off between encapsulation ability and reproductive effort in decorated crickets. Am Nat 2018;191:452–74. doi:10.1086/696147.

[34] Dinh H, Mendez V, Tabrizi ST, Ponton F. Macronutrients and infection in fruit flies. Insect Biochem Mol Biol 2019;110:98–104. doi:10.1016/j.ibmb.2019.05.002.

[35] Moadeli T, Taylor PW, Ponton F. High productivity gel diets for rearing of Queensland fruit fly, Bactrocera tryoni. J Pest Sci (2004) 2017;90:507–20. doi:10.1007/s10340-016-0813-0.

[36] Nguyen B, Ponton F, Than A, Taylor PW, Chapman T, Morimoto J. Interactions between ecological factors in the developmental environment modulate pupal and adult traits in a polyphagous fly. Ecol Evol 2019;9:ece3.5206. doi:10.1002/ece3.5206.

[37] R Development Core Team, 2011. R: A Language and environment for statistical computing. Vienna, Austria: the R Foundation for Statistical Computing. ISBN: 3-900051-07-0. http://www.R-project.org/.

[38] Cressler CE, Nelson WA, Day T, Mccauley E. Disentangling the interaction among host resources, the immune system and pathogens. Ecol Lett 2014;17:284–93. doi:10.1111/ele.12229.

[39] Smith V. Host resource supplies influence the dynamics and outcome of infectious disease. Integr Comp Biol 2007;47:310–6. doi:10.1093/icb/icm006.

[40] Bedhomme S, Agnew P, Sidobre C, Michalakis Y. Virulence reaction norms across a food gradient. Proc R Soc B Biol Sci 2004;271:739–44. doi:10.1098/rspb.2003.2657.

[41] Pulkkinen K, Ebert D. Host starvation decreases parasite load and mean host size in experimental populations. Ecology 2004;85:823–33. doi:10.1890/03-0185.

[42] Hall SR, Simonis JL, Nisbet RM, Tessier AJ, Cáceres CE. Resource ecology of virulence in a planktonic host-parasite system: An explanation using dynamic energy budgets. Am Nat 2009;174:149–62. doi:10.1086/600086.

[43] Muturi EJ, Kim CH, Alto BW, Berenbaum MR, Schuler MA. Larval environmental stress alters Aedes aegypti competence for Sindbis virus. Trop Med Int Heal 2011;16:955–64. doi:10.1111/j.1365-3156.2011.02796.x.

[44] Duneau D, Ferdy JB, Revah J, Kondolf H, Ortiz GA, Lazzaro BP, et al. Stochastic variation in the initial phase of bacterial infection predicts the probability of survival in D. melanogaster. Elife 2017;6. doi:10.7554/eLife.28298.

[45] Ponton F, Morimoto J, Robinson K, et al. Macronutrients modulate survival to infection and immunity in Drosophila. J Anim Ecol 2020; 89:460–470. doi.org/10.1111/1365-2656.13126.

[46] Boots M, Begon M. Trade-offs with resistance to a Granulosis virus in the Indian meal moth, examined by a laboratory evolution experiment. Funct Ecol 1993;7:528. doi:10.2307/2390128.

[47] Rantala MJ, Roff DA. An analysis of trade-offs in immune function, body size and development time in the Mediterranean field cricket, Gryllus bimaculatus. Funct Ecol 2005;19:323–30. doi:10.1111/j.1365-2435.2005.00979.x.

[48] Stefana MI, Driscoll PC, Obata F, Raquel Pengelly A, Newell CL, MacRae JI, Gould AP. Developmental diet regulates Drosophila lifespan via lipid autotoxins. Nature Comm 2017;8(1): 1384.

[49] Hoshizaki, D. (2012). Fat body. In R. Chapman (Author) & S. Simpson & A. Douglas (Eds.), The insects: Structure and function (pp. 132–146). Cambridge: Cambridge University Press. doi:10.1017/CBO9781139035460.009 (ref no. 46)

[50] Langenhof MR, Komdeur J. Why and how the early-life environment affects development of coping behaviours. Behav Ecol Sociobiol 2018;72:34. doi:10.1007/s00265-018-2452-3.

[51] Boots M, Roberts KE. Maternal effects in disease resistance: Poor maternal environment increases offspring resistance to an insect virus. Proc R Soc B Biol Sci 2012;279:4009–14. doi:10.1098/rspb.2012.1073.

[52] Ben-Ami F, Ebert D, Regoes RR. Pathogen dose infectivity curves as a method to analyze the distribution of host susceptibility: A quantitative assessment of maternal effects after food stress and pathogen exposure. Am Nat 2009;175:106–15. doi:10.1086/648672.

[53] Mitchell SE, Read AF. Poor maternal environment enhances offspring disease resistance in an invertebrate. Proc R Soc B Biol Sci 2005;272:2601–7. doi:10.1098/rspb.2005.3253.

[54] Moreno-Garcïa M, Lanz-Mendoza H, Córdoba-Aguilar A. Genetic variance and genotype-by-environment interaction of immune response in Aedes aegypti (Diptera: Culicidae). J Med Entomol 2010;47:111–20. doi:10.1603/me08267.

[55] Rosa E, Saastamoinen M. Sex-dependent effects of larval food stress on adult performance under semi-natural conditions: only a matter of size? Oecologia 2017;184:633–42. doi:10.1007/s00442-017-3903-7.

[56] Duxbury EML, Chapman T. Sex-specific responses of lifespan and fitness to variation in developmental versus adult diets in D. melanogaster. Journals Gerontol Ser A 2020;75:1431–1438. doi:10.1093/gerona/glz175.

[57] Despland E, Noseworthy M. How well do specialist feeders regulate nutrient intake? Evidence from a gregarious tree-feeding caterpillar. J Exp Biol 2006;209:1301–9. doi:10.1242/jeb.02130.

[58] Sim C, Denlinger DL. Insulin signaling and the regulation of insect diapause. Front Physiol 2013;4:189. doi:10.3389/fphys.2013.00189.

[59] Mizoguchi A, Okamoto N. Insulin-like and IGF-like peptides in the silkmoth Bombyx mori: Discovery, structure, secretion, and function. Front Physiol 2013;4 AUG. doi:10.3389/fphys.2013.00217.

[60] Nässel DR, Liu Y, Luo J. Insulin/IGF signaling and its regulation in Drosophila. Gen Comp Endocrinol 2015;221:255–66. doi:10.1016/j.ygcen.2014.11.021.

[61] Zoncu R, Bar-Peled L, Efeyan A, Wang S, Sancak Y, Sabatini DM. mTORC1 senses lysosomal amino acids through an inside-out mechanism that requires the vacuolar H+-ATPase. Science (80-) 2011;334:678–83. doi:10.1126/science.1207056.

[62] Chantranupong L, Wolfson RL, Sabatini DM. Nutrient-sensing mechanisms across evolution. Cell 2015;161:67–83. doi:10.1016/j.cell.2015.02.041.

[63] Fu K-Y, Guo W-C, Ahmat T, Li G-Q. Knockdown of a nutrient amino acid transporter gene LdNAT1 reduces free neutral amino acid contents and impairs Leptinotarsa decemlineata pupation. Sci Rep 2015;5:18124. doi:10.1038/srep18124.

[64] Sutherland WJ, Freckleton RP, Godfray HCJ, Beissinger SR, Benton T, Cameron DD, et al. Identification of 100 fundamental ecological questions. J Ecol 2013;101:58–67. doi:10.1111/1365-2745.12025.

[65] Acevedo-Whitehouse K, Duffus ALJ. Effects of environmental change on wildlife health. Philos Trans R Soc B Biol Sci 2009;364:3429–38. doi:10.1098/rstb.2009.0128.

[66] Deem SL, Karesh WB, Weisman W. Putting theory into practice: Wildlife health in conservation. Conserv Biol 2001;15:1224–33. doi:10.1111/j.1523-1739.2001.00336.x.

